# Antiferromagnetic switch in serum

**DOI:** 10.1101/211102

**Authors:** Sufi O. Raja, Sanjay Chatterjee, Anjan Kr. Dasgupta

## Abstract

1.

Ferritin contains naturally occurring iron oxide nanoparticle surrounded by a structured spherical array of peptide residues that provides tremendous stability to this iron storage protein. We use synthetic citrate coated Super Paramagnetic Iron Oxide Nanoparticles (SPIONs) and static magnetic field in exploring the Ferritin induced magnetic environment of human serum samples with varying ferritin level collected from thalassemic patients. We report anti-ferromagnetic properties of serum in patients with iron overloading. Magnetic pulling by an external magnetic field showed a cusp-like behavior with increasing concentration of serum Ferritin measured by standard ELISA based kit. A reduction in the extent of pulling after a threshold concentration of Ferritin (1500 ng/ml) suggests a Ferritin dependent magnetic switching.Negative magnetization (anti-ferromagnetization) was confirmed by Vibrating Sample Magnetometric (VSM) analysis of SPION-serum mixture containing very high level of Ferritin. Such magnetic switching may have a possible role in iron homeostasis during overloading of Ferritin.

**Abbreviations:** SPIONs: Super Paramagnetic Iron Oxide Nanoparticles, VSM: Vibrating Sample Magnetometry, SQUID: Super conducting Quantum Interference Device, PCS: Photon Correlation Spectroscopy

## 4. Introduction

Ferritin is a non-heme iron containing protein which plays crucial roles in body iron homeostasis^1^. In general serum Ferritin level is considered as “*low cost*” reliable marker ^2^ for body iron homeostasis. But, several case studies have pointed out that only serum Ferritin measurement is not sufficient to understand the body iron management scenario of an individual as modulation of serum Ferritin is also associated with diverse physiological and pathologic processes^3–5^. Briefly, apart from iron related disorders^6^, modulation of serum Ferritin is associated with chronic inflammation^7,8^ and neurological disorders ^9,10^. Hence, clinicians frequently face problems to identify the clinical status whether it is purely iron related disorders (except genetic disorders like Hereditary hemochromatoses, a genetic disorder causes iron overloading^11^ or not. As the biopsy of liver tissue and its iron staining (gold standard) is an invasive method, clinicians generally recommend some non-invasive techniques to overcome aforementioned problems^12^. Some of them are measurement of soluble transferrrin receptor (sTfR)^13^, transferrin saturation (TSAT)^14^ and reticulocyte hemoglobin content^15^. In general, some *empirical* thresholds have been set to identify clinical symptoms associated with altered serum Ferritin level.From molecular point of view attempts have been made to explore the nature of Ferritin during various clinical symptoms. But, no useful variation in their structure has been noticed. One important finding was alteration of degree of glycosylation in serum Ferritin during various pathological conditions^16^. But, exploring the clinical significance of such difference needs further research.

Apart from the clinical and physiological significance of Ferritin, it is also an interesting candidate in material science due to occurrence of emerging properties at nanoscale. Ferritin is a naturally occurring iron oxide nanoparticles^17^. Ferritin sequesters iron (Fe^+2^) and mineralize it within its core (~8 nm diameter) which is coated by polypeptide shell^18^. Magnetic properties of purified Ferritin from horse spleen have been extensively studied. It resembles several exotic magnetic properties like super-paramagnetism that is found in synthetic iron oxide nanoparticles ^19,20^. It is also magnetically birefrigent^21^. Instead of such knowledge, physiological role of nanoscale magnetism of Ferritin in body iron homeostasis is not well explored. The possible role was pointed out several years ago by Hilty and coworkers^22^, where they showed that magnetic property of Ferritin is highly dynamic and depends on iron loading within the core. But still we do not have clear understanding about the functional correlation between magnetic property of Ferritin and iron homeostasis. Here we have tried to explore such correlation in the context of iron regulation during thalassemia. In addition we have also developed a low cost alternative qualitative technique of whole body SQUID, which is most preferable technique by clinicians to get true snapshot of body iron management. The assay is based on probing the magnetic interaction between synthetic iron oxide nanoparticles and human serum by external magnetic field, which. The use of magnetic nanoparticles in probing the magnetic environment of serum in the proposed way is a new and novel approach to the best of our knowledge.

## 5. Materials and Methods

### 5.1. Synthesis of Iron Oxide nanoparticles

The detail of the synthesis protocol is described elsewhere^23^. Briefly, we have prepared iron oxide nanoparticles by co-precipitating 2gm ferrous chloride and 4gm ferric chloride (solubilized in 50 ml 2(N) HCl) by 100ml 1.5(N) sodium hydroxide upon constant stirring at room temperature. The precipitate was washed well with milli-Q water through magnetic decantation to remove the excess/un-reacted ionic species. Then 20ml Citrate buffer (1.6gm citric acid and 0.8gm tri-sodium citrate) was added to collect the stabilized nanoform. Finally pH was adjusted to ~7 using 1.5M NaOH. Then stable colloidal suspension was centrifuged at 10,000 rpm for 30 minutes to get rid of the larger aggregates and sup was stored at 4 degree Celsius. Concentration of nanoparticles solution in terms of ionic iron was measured by Ferrozine assay.

### 5.2. Preparation of blood serum

Blood samples were collected from individuals after obtaining informed consent patients admitted to Institute for Hematology and Transfusion Medicine (IHTM), Kolkata with normal and elevated level of serum ferritin.. The study protocol was approved by the Institutional Ethical Committee of IHTM.

### 5.3. Interaction of Nanoparticles with human blood serum

Serum samples are centrifuged at 2000 rpm for 10 minutes and the supernatant was taken as working serum solution. In a 0.5ml centrifuge tube, 10 µl nanoparticle solutions was added to 100 µl of serum. They were incubated both in absence and in presence of a strong bar magnet (90 mTesla). After incubation, the supernatant for both the sets was checked spectrophotometrically (Thermo-Fisher). Absorbance was measured at 280 nm to measure the protein as well as nanoparticles concentration in different samples. The difference in absorbance in presence and absence of magnetic field was calculated for the measurement of percent decrease of this absorbance with respect to initial absorbance at the said wavelength.

### 5.4. Size measurement

The hydrodynamic size and zeta potential were measured by Photon Correlation Spectroscopy (Malvern Nano-ZS) using the dynamic light scattering property.

### 5.5. Ferritin assay

Ferritin assay of the serum samples were done using ORG 5FE kit (ORGENTIC Diagonisticka GmbH). Anti-human-ferritin antibodies are bound to microwells. Ferritin, present in diluted serum, or plasma bind to the respective antibody. Washing of the microwells removes unspecific serum and plasma components. Horseradish peroxidase (HRP) conjugated anti-human ferritin immunologically detects the bound patient ferritin forming a conjugate/ferritin/antibody complex. Washing of the microwells removes unbound conjugate. An enzyme substrate in the presence of bound conjugate hydrolyses to form a blue color. The addition of an acid stops the reaction forming a yellow end-product. The intensity of this yellow color is measured photometrically at 450nm. The amount of color is directly proportional to the concentration of ferritin present in the original sample.

### 5.6. Characterization of magnetic properties of iron oxide nanoparticles

The magnetic nanoparticles were characterized by Superconducting Quantum Interference Device (SQUID) [MPMS-7 (Quantum Design)]. For the zero field cooled (ZFC) data, the sample was cooled down to 5 K in the absence of magnetic field and the data were taken while warming up to 300 K in the presence of 100 Oe field. The field cooled (FC) data were taken while warming the sample after cooling it to 5 K in the presence of the 100 Oe field. The hysteresis data were taken at various temperatures up to 300 K and up to a field of 4 T (40000 Oe).

## 6. Results and Discussion

Dynamic light scattering studies confirm nearly mono-dispersed (polydispersity index ~ 0.27) population of synthesized citrate coated iron oxide nanoparticles with mean size of ~20 nm (see Figure S1). Atomic force microscopic analysis and surface conjugation of this nanoparticle was shown previously by us^23^. Here we performed detailed magnetic analysis of iron oxide nanoparticles using Vibrational Sample Magnetometer (VSM). Generally, iron oxide nanoparticles beyond a certain size regime show an emergent magnetic property-super paramagnetism. The existence of super-paramagnetic behavior of such particles is studied through magnetization measurement in absence (zero filed cooling, ZFC) and in presence of 100 Oe external magnetic field (field cooling, FC), respectively. The distinct bifurcation of the ZFC and FC plot around 80 K (see Figure 1a) clearly indicates blocking temperature (T_B_) as indicated by an arrow in the figure. The ZFC magnetization increases as a function of temperature for temperature 5K<T<80 K, indicating that there are blocked moments which start to contribute to the magnetization when the temperature is increased. In the same temperature region the FC magnetization decreases as the temperature increases, since the field induced aligned moments start randomizing due to thermal energy. Beyond the blocking temperature both the ZFC and FC curve coincide and the magnetization decreases as the temperature increases due to temperature induced randomization of the moments. This behavior is typical of super paramagnetic nanoparticles and is shown in Figure 1a.

**Figure 1:**
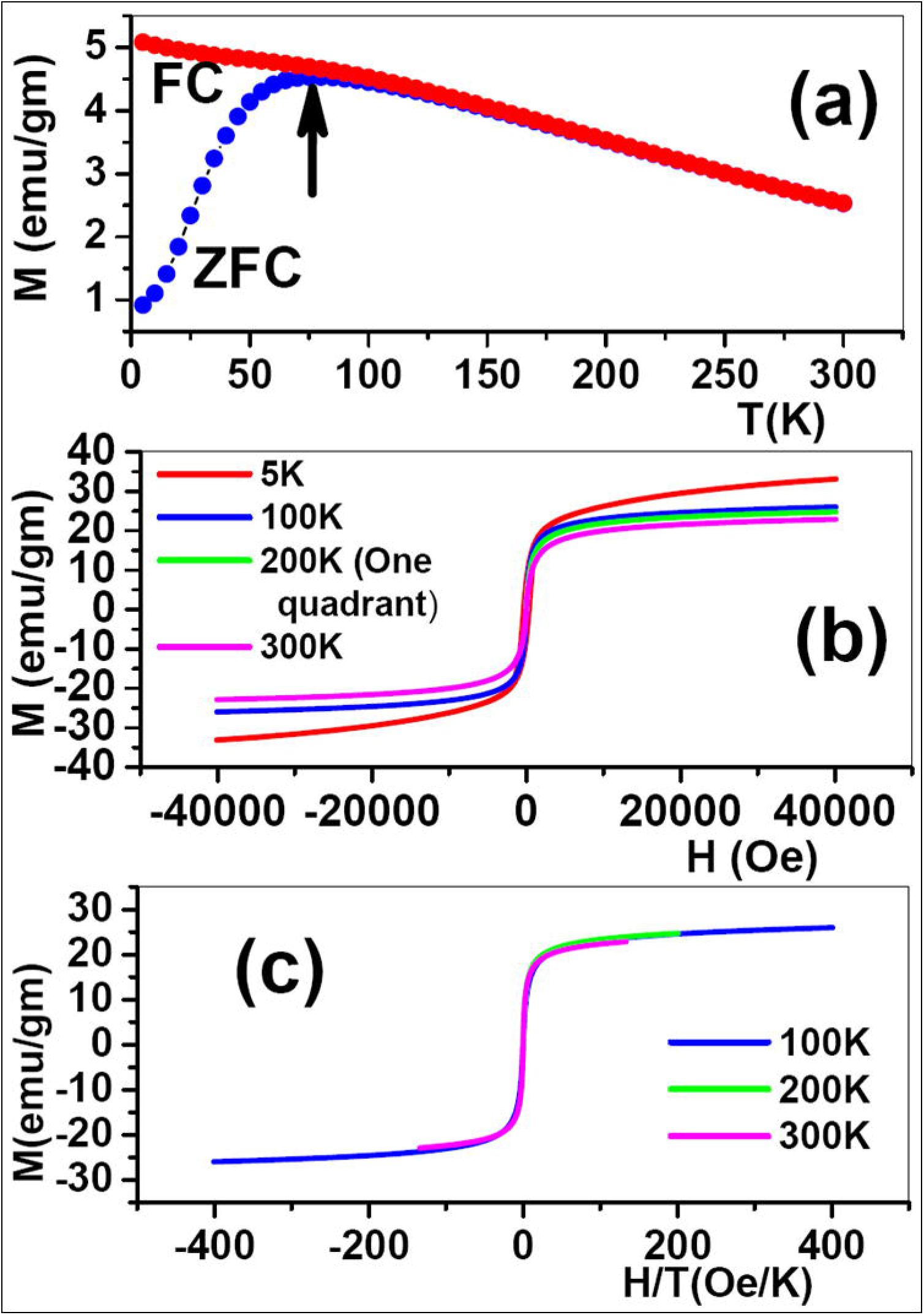
SQUID based magnetic characterization of synthetic citrate coated iron oxide nanoparticles.**(a)**Zero Field Cooling (ZFC) and Field Cooling (FC) magnetization curves taken at 100 Oe. Blocking temperature (T_B_) is shown by the arrow. **(b)** Magnetization (M) versus applied magnetic field (H) plot of the sample taken at 5K, 100K, 200K and 300K up to 4T. **(c)** Magnetization data OF (b) re-plotted as a function of H/T.

In Figure 1b we show variation of magnetization (M) versus applied magnetic field (H) of the sample taken at temperature of 5K, 100K, 200K and 300K up to 4 Tesla. We observed “no hysteresis” for 100K, 200K and 300K data (above T_B_) and the M-H curve shows S-shaped behavior with saturation similar to a superparamagnetic behavior. We also observed no saturation in the M-H curve taken at 5 K up to 4T. To verify that the iron oxide nanoparticles are ideal superparamagnetic, one should observe that the magnetization isotherms scale with H/T apart from no hysteresis. We have re-plotted the magnetization from Figure 1b in Figure 1c as a function of H/T and we indeed observe the that the magnetization scales with H/T and all the three curves for temperature above T_B_ (100K, 200K and 300K) collapses on one single master curve as predicted for superparamagnetic nanoparticles, indicating that each of the particles are within single domain regime.

Next we studied the interaction of the synthesized super paramagnetic iron oxide nanoparticles (SPIONs) with human serum which contains thousands of protein including iron storage protein ferritin. Ferritin is also a naturally occurring iron oxide nanoparticle. Hence, we were interested to study the interaction between natural and synthetic magnetic nanoforms. There is an existing report where such interaction was studied using *purified* Ferritin^24^. But, the present study, to the best of our knowledge is the first attempt where we have tried to probe the Ferritin induced magnetic environment of human serum using SPIONs. As thalassemic patients have higher level of serum Ferritin due to frequent transfusion of blood, we collected serum samples from thalassemic patients with varying serum Ferritin level. To probe the interaction between SPIONs and serum we incubated them in absence and presence of magnet (70 mTesla) as shown in Figure 2a. As expected magnetic field results in pull down of some fraction. To quantify the extent of magnetic pulling we measured the absorbance at 280 nm of the supernatant with or without magnetic incubation. Then we plotted the % change of absorbance (% ΔOD) with serum ferritin concentration measured by ELISA based kit. Surprisingly we found a cusp like behavior as shown in the Figure 2b.The % decrease in absorbance at 280 nm shows a prominent criticality. When the ferritin level reaches the range 1400-1500 ng/ml, the extent of magnetic pulling is drastically reduced. This anomaly is however well explained on the light of anti-ferromagnetism. Studies on reconstituted Ferritin showed the occurrence of antiferromagnetic exchange interaction between iron atoms upon increase in number of iron atoms within ferritin’s core. Hence, it is evident from our study also that the magnetic property of Ferritin is modulated by level of Ferritin in the serum. The Ferritin, when it is very high in the serum interacts with SPION differently and results in reduced affinity towards magnetic field.

**Figure 2:**
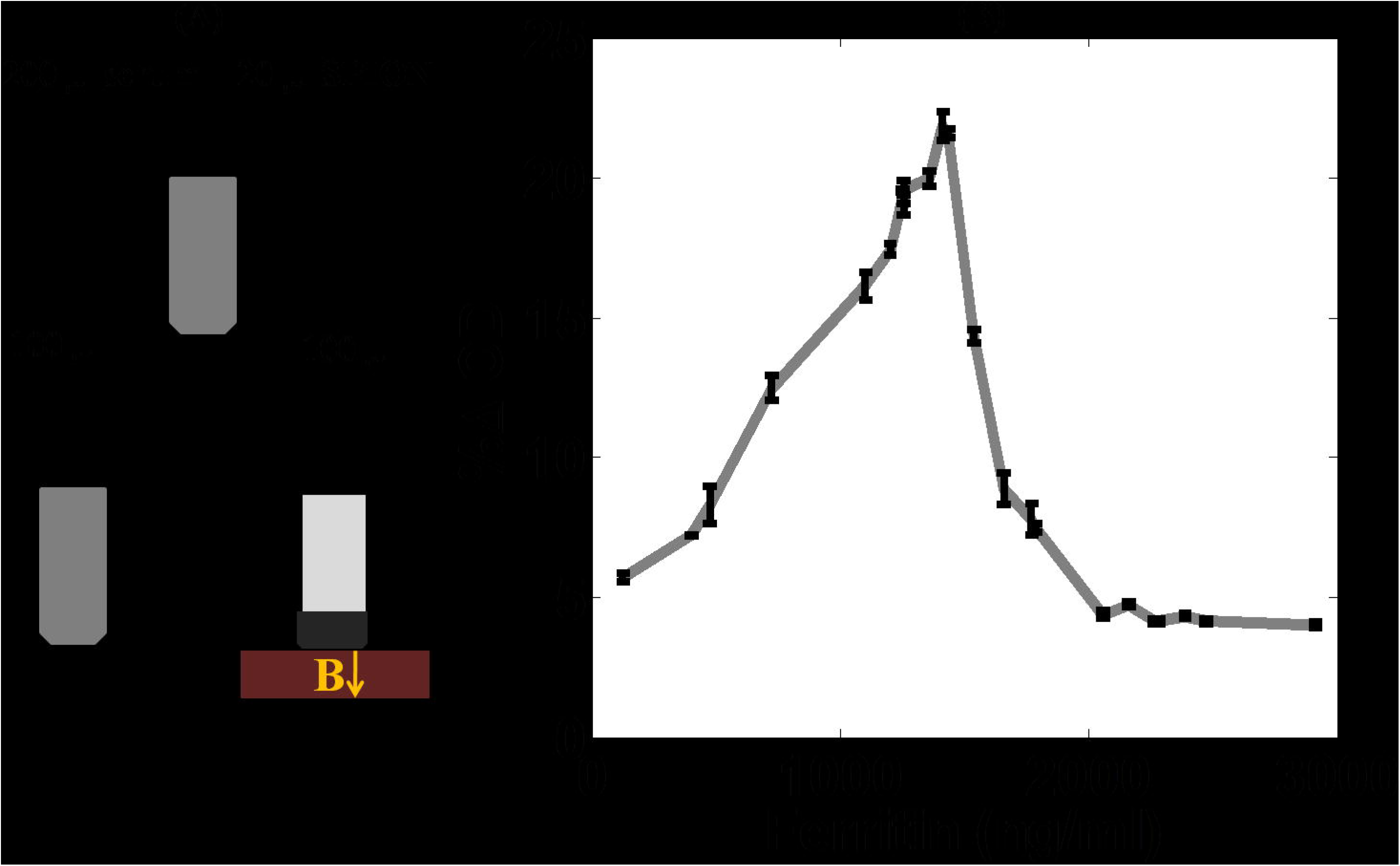
(a) shows the schematic description of the proposed magnetic pulling assay. After the assay absorbance was measured for the supernatants and the percentage difference (%∆ OD, Y-axis) were plotted against serum ferritin level (X-axis) measured by standard ELISA assay (b). A cusp like dependence of %∆ OD on serum ferritin concentration indicates loss of magnetic affinity of serum-SPIONs mixture after a certain threshold concentration of serum ferritin.

Next we wanted to check whether the effect is due to increased concentration of Ferritin protein or due to change in magnetic property of feritin’s core. So, we performed the same magnetic pulling assay using pure horse spleen Ferritin and took bovine serum albumin as control. But, we did not find any significant change in %Δ OD with increasing Ferritin concentration. The % Δ OD was more or less same for Ferritin and BSA. We believe that this is not the proper control as the overall protein content of serum is several folds higher. Hence, there must be difference in colloidal stability. Few nanograms level of purified protein in 100 mM phosphate buffer basically results in colloidal instability of the nanoaprticle. That is why we found sedimentation for both the cases (with or without magnetic field) which is the reason of observed low value for %Δ OD. But in serum, nanoparticles can withstand the high ionic stress due to presence of plenty of proteins. Hence the observed pulling in presence of magnet is due to pure magnetic interaction. In this case it is unlikely that only nanogram level of Ferritin is bound to the surface of the particle. Rather non-specific interaction of several other proteins is a must for getting colloidal stability in a medium of high ionic strength like serum. But, it is logical to propose that presence of Ferritin of different magnetic property within the protein assembly on the surface of SPION may result in differential affinity towards magnetic field. There are several reports exists including ours on modulation of effective magnetic property upon surface modification.

We also performed ELISA based experiments after the magnetic incubation step to check the amount of Ferritin in the pellet. Figure 3 shows 450 nm OD for the samples with increasing serum Ferritin level for the serum only (blue), serum and SPION mixture with (red) or without (green) magnetic incubation respectively. The observed increased value of OD for serum-SPIONs mixture compared to only serum is due to contribution from intrinsic peroxidase activity of SPION. SPIONs intrinsic activity adds on to the horse radish peroxidase activity which is used as secondary antibody in the ELISA based assay. Attachment of SPION on the ELISA plate directly proves the association of serum Ferritin with SPIONs. Now if this attachment was a random event then we should observe random trend of OD with increasing serum Ferritin level and if the attachment was independent of nature of magnetic interaction between serum Ferritin and SPION then we should always see increased value of OD for the sample with magnetic incubation than without incubation irrespective of the serum Ferritin level. But, here also we observed that after a certain concentration of Ferritin the OD value for un-magnetized sample is greater than the magnetized sample. It suggests high Ferritin level is somehow results in lower affinity of the serum-SPIONs mixture towards magnet.

**Figure 3:**
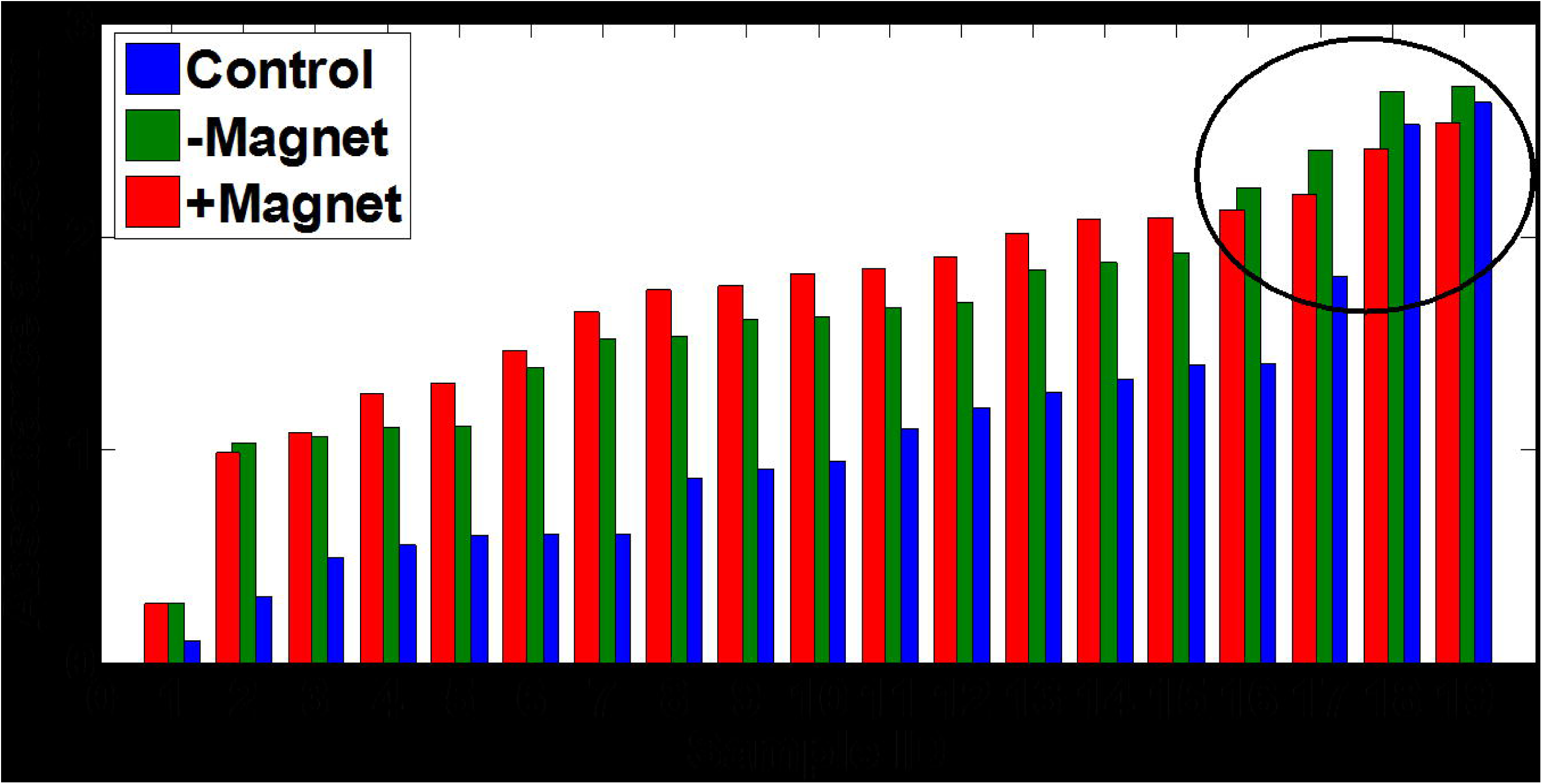
After the magnetic pulling assay pellets were transferred to ELISA plate. ELISA assay was performed along with the crude serum (without SPIONs) to check the serum ferritin level. Bar represents the raw ELISA reading at 450 nm where blue, green and red stand for control serum and serum-SPION mixture with or without magnetic incubation respectively. Here also like Figure 2 we got reduction in extent of magnetic pulling after a threshold concentration of serum ferritin. Serum ferritin level of the samples is given in the supplementary section.

Finally, to probe whether high serum Ferritin is actually responsible for differential magnetic environment of serum or not we performed VSM analysis of the SPION-serum mixture containing low varying concentration of serum Ferritin (100.2, 1069.2 and 2139.1 ng/ml). Figure 4 clearly shows reduction of magnetization for the SPION-serum mixture containing high Ferritin (2139.1 ng/ml). We found reduced magnetic susceptibility for this sample (see Figure 4b) as well as lowest value of maximum magnetization below and above T_B_ (see Figure 4c and 4d). Figure 4c clearly shows that the T_B_ values for all the samples are more or less same which indicates that the effective magnetic volume is identical for all these samples. Inspite of having same magnetic volume the reduction in magnetization suggests the occurrence of negative exchange interaction. The source of occurrence of negative exchange interaction may be due to various reasons, one being the canted anti-ferromagnetism^25^. The negative magnetization during ZFC is suggestive of such negative exchange interaction. While the molecular basis of such magnetic switch in our system is yet to explore, our observation indicates the possible functional role in body iron management of such ferritin level dependent differential magnetization of serum.

**Figure 4:**
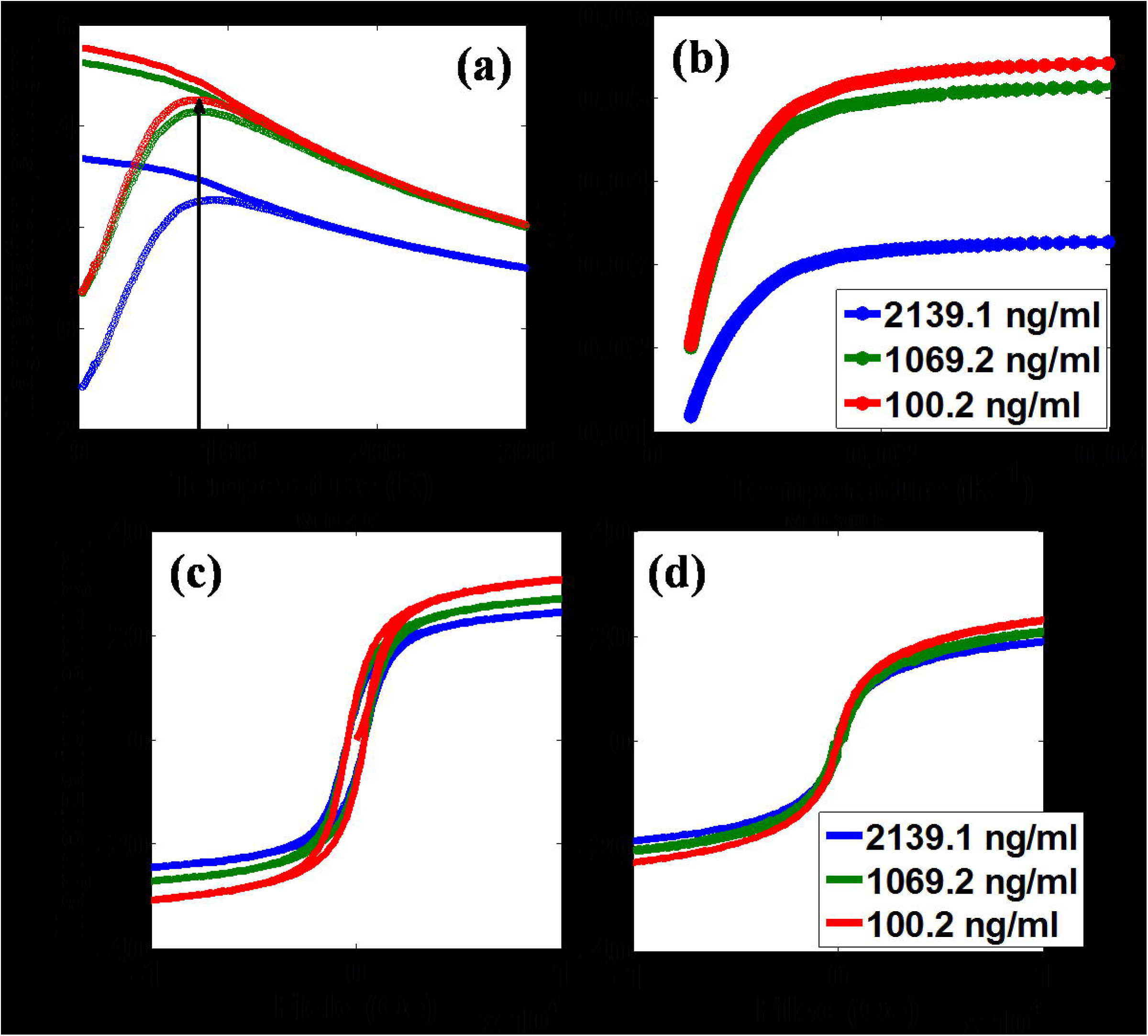
Magnetic measurement of serum-SPIONs mixture. (a) FC-ZFC plots for serum-SPIONs mixture containing different level of ferritin. (b) Shows the magnetic susceptibility of the same samples and (c and d) show the M-H plot at 4K and 300 K. All these plots indicate a reduction of net magnetization for the sample containing highest serum ferritin.

Lastly, from application point of view this simple assay can be an easy and cheap technique compared to commercially available ELISA based technique for serum Ferritin measurement. Up to ~1500 ng/ml we found nearly linear correlation between ΔOD and serum Ferritin. So, one can use this assay to measure serum Ferritin (we patented the technique)^26^. But, how one can differentiate that a low value of ΔOD is due to low or very high serum Ferritin. To resolve this issue we performed a simple assay which also supports the antiferromagnetic exchange interaction based reduction in magnetization. We performed magnetic pulling experiment for the diluted serum containing low and very high Ferritin. We observed decrease in ΔOD with dilution for the serum sample containing low Ferritin (see Figure 5A). Interestingly we got enhancement of ΔOD with dilution for the serum sample containing very high Ferritin (see Figure 5B). It is obvious if we reduce the source which is responsible for reduced magnetization that will definitely increase the extent of magnetic pulling. This observation also suggests that high Ferritin çing serum actually introduce antiferromagnetic interaction with SPION.

**Figure 5:**
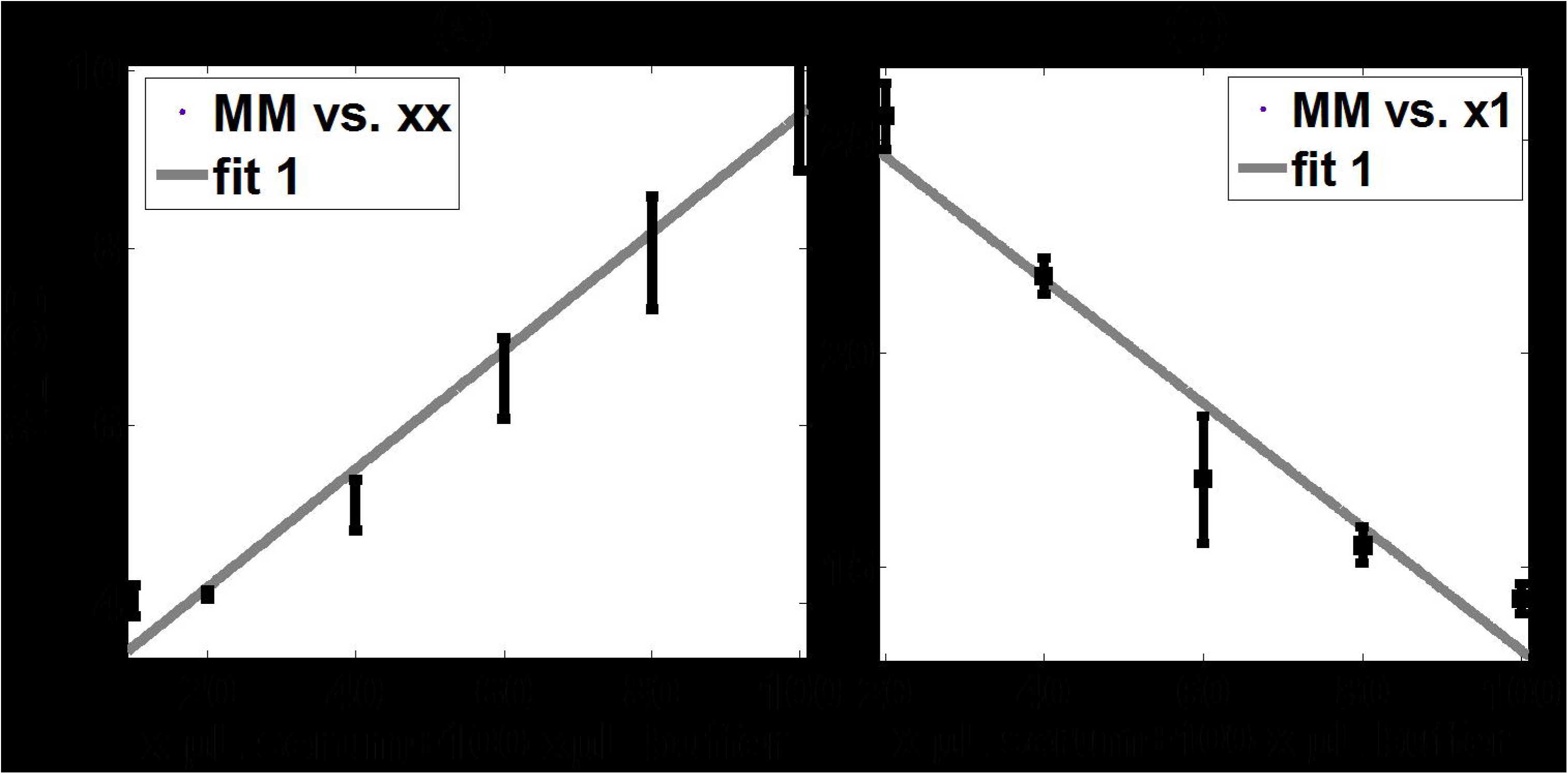
Dilution experiment to resolve the degenerate value of %∆ OD. (a & b) show the effect of dilution with 100 mM phosphate buffer on %∆ OD. Low ferritin (a) containing serum sample shows decrease in %∆ OD with dilution whereas high ferritin (b) containing serum sample shows increase in %∆ OD with dilution.

## 7. Conclusion

In this paper a new approach for quantifying interactions between magnetic nanoparticles and serum in presence and absence of magnetic field has been described. The variations of such interactions with changes in the ferritin level has been observed. The approach provides important insights in anti-ferromagnetism in dictating the interactions between nanoparticles and serum ferritin. The method used may be exploited as a Elisa independent ferritin kit and clinical assessment for patients with Irion overloading can be better assessed.

## 8. Acknowledgements

The authors acknowledge research support from ICMR (5/3/8/322/2016-ITR) and fellowship support from DST-SERB. The authors also thank Prof. S. Kundu and Azhardduin, University of Calcutta for providing access and technical assistance in SQUID measurement.

